# Evidence for distinct isotopic composition of sap and tissue water in tree stems: consequences for plant water source identification

**DOI:** 10.1101/2020.06.18.160002

**Authors:** Adrià Barbeta, Régis Burlett, Paula Martín-Gómez, Bastien Fréjaville, Nicolas Devert, Lisa Wingate, Jean-Christophe Domec, Jérôme Ogée

**Author notes:** Corresponding author name(s): Adrià Barbeta and Jérôme Ogée. and. Author Contributions A.B., J.O. and R.B. designed research; A.B. and N.D. performed field sampling; R.B., A.B. and P.M. performed sap water extractions; B.F. performed bulk stem water extractions; N.D., P.M., J.C.D, J.O. and L.W. performed stable isotope analyses; A.B. and J.O performed the data analysis and wrote the manuscript, with contributions from all authors.

## Abstract

For decades, theory has upheld that plants do not fractionate water isotopes as they move across the soil-root interface or along plant stems. This theory is now being challenged by several recent studies reporting that the water held in woody stems has an isotopic composition that cannot be attributed to any potential water source. Isotopic offsets between stem and source water still need to be explained, as they prevent identifying unambiguously tree water’s origin from water isotope measurements. Here we show that isotopic offsets between stem and source water can be explained by micrometer-scale water isotope heterogeneity within woody stems and soil micropores. Using a novel technique to extract sap water in xylem conduits separately from the water held in other xylem tissues, we show that these non-conductive xylem tissues are more depleted in deuterium than sap water. We also report that, in cut stems and well-watered potted plants, the isotopic composition of sap water reflects well that of irrigation water, demonstrating that no isotopic fractionation occurs during root water uptake or the sap water extraction process. Previous studies showed that isotopic heterogeneity also exists in soils at the pore scale where water adsorbed onto soil particles is more depleted than capillary/mobile soil water. Data collected at a beech (*Fagus sylvatica*) forest indicate that sap water matches best the capillary/mobile soil water from deep soil horizons, indicating that micrometer-scale water isotope heterogeneity in soils and stems must be accounted for to unambiguously identify where trees obtain their water within catchments.

**Significance Statement:** Forests are prime regulators of the water cycle over land. They return, via transpiration, a large fraction of precipitation back to the atmosphere, influence surface runoff, groundwater recharge or stream flow, and enhance the recycling of atmospheric moisture inland from the ocean. The isotopic composition of water in woody stems can provide unique information on the role forests play in the water cycle only if it can be unambiguously related to the isotopic composition of source water. Here, we report a previously overlooked isotopic fractionation of stem water whereby non-conductive tissues are more depleted in deuterium than sap water, and propose a new technique to extract sap water separately from bulk stem water to unambiguously identify plant water sources.

## Introduction

### Bulk stem water is not only sap water

In the xylem of woody plants, sap water flows from roots to leaves through the apoplastic network of vessels or tracheids (Pickard, 1981). Based on early evidence in hydroponic systems that no isotopic fractionation occurs during root water uptake (Washburn & Smith, 1934; Zimmermann *et al*., 1967), the isotopic composition of bulk stem water is often considered to reflect that of plant source water, at least in plants with suberized stems that prevent bulk stem water evaporative loss and isotopic enrichment (Dawson & Ehleringer, 1993). However bulk stem water is not only sap water. Living parenchyma and phloem cells (‘symplastic’ water), as well as the intercellular spaces between xylem cells (‘capillary’ water) also contain water, and potentially provide some to the xylem conduits (Tyree and Yang 1990; Jupa et al. 2016). Additionally, a relatively large amount of water is also present within cell walls (‘fiber’ water), bound via hydrogen bonds to cellulose, hemi-cellulose and, to a lesser extent, lignin (Berry & Roderick, 2005).

### Bulk stem water is not a good indicator of plant source water

Current techniques to collect water from woody stems do not allow the separation of these four different water pools. Most commonly water is extracted from excised stems placed in a vacuum line and heated to release water vapor into the line where after it is collected in a cryogenic trap. With the conditions of pressure (<10 Pa) and temperature (typically 60-80°C) used, this technique collects all of the stem water pools. There is however growing evidence that total (bulk) stem water collected this way is not a good indicator of plant source water. If this was the case, the isotopic composition of bulk stem water should reflect a weighted mean of the isotopic composition of plant water sources, most often soil water from different horizons. However, the isotopic composition of bulk stem water is more often depleted in ^2^H than any potential water source (Zhao *et al*., 2016; Barbeta *et al*., 2019, 2020; Poca *et al*., 2019). This is illustrated in Figs. 1a-b where bulk stem water from a riparian *F. sylvatica* forest displays δ^2^H values that are more negative than the δ^2^H values of soil water from top and deep horizons, or ground and river water. Similar offsets from the same riparian forest have been reported previously for an entire growing season and also shown to occur in another species (*Quercus robur*) and trees of different stature (Barbeta *et al*., 2019). More recently, we have shown that such δ^2^H offsets between bulk stem and soil water can also be found in potted *F. sylvatica* saplings grown in semi-controlled glasshouse environments (Barbeta *et al*., 2020).

**Figure 1.**
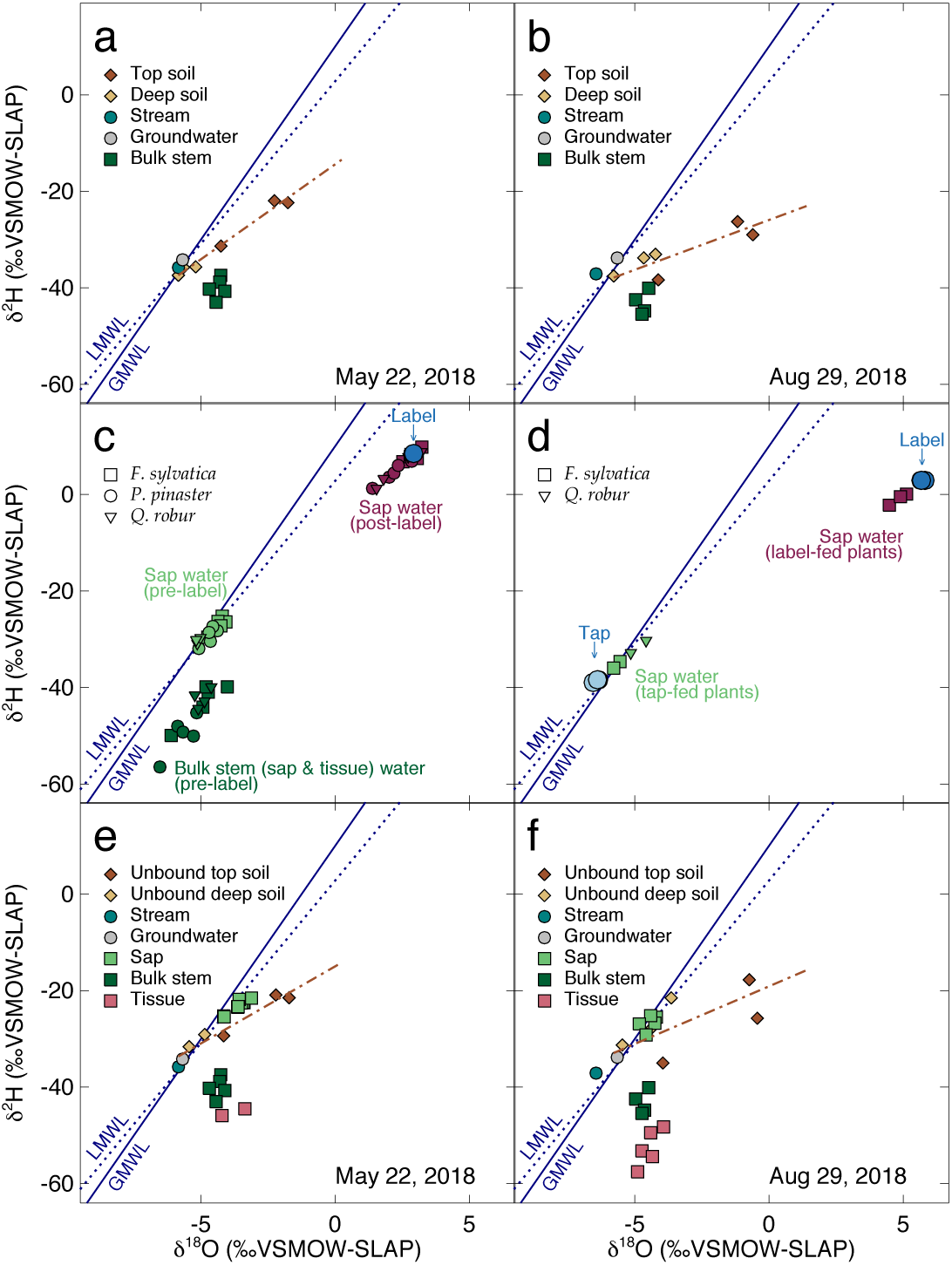
Dual isotope representation of different water pools. Panels (a-b) show the isotopic separation between bulk stem water from mature *Fagus sylvatica* branches and potential water sources (top and deep soil, groundwater and stream water) sampled at two dates during the growing season at the study site, a riparian forest in the SouthWest of France. Panel (c) shows the isotopic separation between bulk stem water and sap water in cut branches sampled from the same field site and the good agreement of sap water with meteoric water (in pre-labelled branches from the field) and with label water (in the same branches after flushing them with label water). Panel (d) shows the good agreement of irrigation water with sap water extracted from the main stem of potted saplings irrigated for several days with either tap or label water. Panels (e-f) show the good agreement of sap water with “unbound” soil water from the same location as in (a-b) (see text for details on how unbound soil water was estimated). LMWL: local meteoric water line. GMWL: global meteoric water line. Brown dot-dashed lines in panels (a-b) and (e-f) indicate the soil evaporation line.

### Initial hypotheses to explain isotopic offsets between soil and stem water

Many other studies have reported isotopic offsets of similar magnitude between soil and stem water, and several hypotheses have been advanced, and often rejected by follow-up studies. Because it was initially thought that such offsets were specific to halophytic and xerophytic plants, a first hypothesis proposed that the suberized root endodermis and developed Casparian strip of these plants fractionated water isotopes during uptake because water was forced to flow through the symplastic route (the continuum of communicating cytoplasms connected by plasmodesmata or aquaporins), expected to cause a more pronounced isotopic fractionation than if water moved through roots via the apoplast (Lin & Sternberg, 1993; Ellsworth & Williams, 2007; Poca *et al*., 2019). This first hypothesis has been challenged by several studies reporting soil-root isotopic offsets in plant species where root water uptake through the apoplastic route should not be impeded (Zhao *et al*., 2016; Vargas *et al*., 2017; Barbeta et al. 2019) and finally rejected by a recent experiment with *F. sylvatica* saplings demonstrating no fractionation by root water uptake despite strong soil-stem water isotope offsets (Barbeta *et al*., 2020).

An alternative hypothesis was proposed in a study of potted *Persea americana* saplings. Because soil-stem isotopic offsets were observed for both ^2^H and ^18^O, and “with a slope close to 8” as for liquid-vapor isotopic equilibration, it was suggested that soil water evaporation, followed by vapor transport and condensation on the root tips may have been responsible for the observed isotopic offsets (Vargas *et al*., 2017). We have argued recently that, as long as the condensed water at the root tip stays in thermodynamic and isotopic equilibrium with water vapor in soil pores, such a chain of reactions cannot create any soil-stem isotopic offset of several per mil amplitude (Barbeta *et al*., 2020). This should be the case in most, if not all, situations, including those in the experiment on *P. Americana* that led to the formulation of the hypothesis. Furthermore, the proposed chain of reactions and associated isotopic fractionations cannot explain why large soil-stem isotopic offsets are found even in well-watered situations where liquid-vapor isotopic equilibration are most expected (Barbeta *et al*., 2020).

### A recent hypothesis to explain isotopic offsets between soil and stem water

Another hypothesis was proposed recently that bulk stem water has an isotopic composition that differs significantly from that of sap (and thus source) water. Support for this hypothesis was obtained by Zhao et al. (2016) who took advantage of the positive root pressure conditions experienced by some riparian tree species to sample sap water using a syringe inserted in the sapwood of such trees. They found that water collected this way had an isotopic composition that differed markedly from bulk stem water but coincided well with that of groundwater, the only plausible water source for these riparian desert trees (Zhao *et al*., 2016). Isotope heterogeneity in stem water pools (i.e. between long-transport conduits and other tissues) was also proposed recently as the most plausible hypothesis to explain isotopic offsets between soil and stem water in potted *F. sylvatica* saplings (Barbeta *et al*., 2020). However, apart from the study by Zhao *et al*. (2016), direct evidence for an isotopic difference between sap water in long-transport conduits and water in other stem tissues is still lacking, and the mechanism behind it is still not known.

### A new technique to extract sap water separately from the water in other stem tissues

Unfortunately, the sap water collection technique used by Zhao and colleagues is only applicable to tree species that experience positive root pressure. Besides, it has not been completely proven that this technique allowed the collection of sap water only. Here, we used a flow-rotor centrifuge originally designed to study the hydraulic embolism resistance of woody stems (Cochard, 2002) and later improved with a custom-made water collector (Pivovaroff *et al*., 2016; Peng et al., 2019). Using staining methods, it was recently shown that this particular centrifuge technique, hereafter called the “cavitron” technique, should mainly extract sap water from xylem conduits (Peng et al., 2019). We therefore applied this sap water extraction technique on cut branches of adult trees over an entire growing season. We also applied the technique to branches that had been refilled with water of known isotopic composition and to stems of potted saplings with known irrigation water.

## Results

### Testing the cavitron technique to extract sap water on cut branches from the field

The water extracted with the “cavitron” technique (denoted ‘sap water’ from hereon) from cut branches of *F. sylvatica, Q. robur* and *P. pinaster* collected from the field (i.e. ‘pre-label’) fell on the local meteoric water line (LMWL; the regression line between δ^18^O and δ^2^H of local rainfall water) (Fig. 1c). Bulk stem water δ^2^H from these ‘pre-label’ branches was more depleted than sap water, by 16‰ on average (*P* < 0.0001) while differences in δ^18^O were not significant (*P* = 0.075). To verify the absence of fractionation during centrifugation we flushed the then-embolized stems with labelled water of a known isotopic composition and re-extracted sap water from the stems with the cavitron technique (see Methods). The δ^18^O and δ^2^H of sap water re-extracted after flushing (i.e. ‘post-label’) was not significantly different from those of the label water (Fig. 1c). Those labelled sap water samples that were slightly isotopically different from the label fell on a mixing line between pre-label sap water values and the labeled water, possibly caused by an incomplete replacement of the former. These results were observed on three tree species and no statistical difference (*P* > 0.05) in the isotopic composition of sap or bulk stem water pools were found amongst the species.

Water from fresh samples was extracted at two different rotation speeds in order to distinguish between water released from open conduits and intercellular spaces (capillary water extracted at less than -2 MPa), and water from intact vessels and tracheids (cavitation water extracted between -2 and -6 MPa when xylem conduits embolized). The threshold of -2 MPa was chosen according to dehydration isotherms previously described for *F. sylvatica* branches (Jupa *et al*., 2016). In some samples, extractions at -2 MPa did not yield enough water for stable isotope analysis. For all the other samples, there were no significant differences in δ^18^O or δ^2^H between water extracted at -2 MPa and -6 MPa (*P* = 0.20 for δ^18^O and *P* = 0.54 for δ^2^H), indicating that the two water pools are well connected.

### Testing the cavitron technique to extract sap water on potted plants

To verify that the water extracted with the cavitron technique was a good indicator of plant source water (and thus sap water) we also performed experiments on potted *F. sylvatica* and *Q. robur* saplings irrigated daily with tap water for 18 days, with the exception of a subset of 3 plants that we switched irrigation from tap to label water for the last 3 days before sampling (see Methods). Results from this experiment showed that, for the two species, the isotope composition of sap water extracted with the cavitron technique (from the main stems of the saplings) matched very closely that of irrigation water, with some small variations (Fig. 1d). In tap-fed plants, the small variations in sap water were aligned along what resembles an evaporation line originating from tap water. In label-fed plants, the narrow distribution of sap water again followed a mixing line between new label water and old tap water (Fig. 1d), consistent with observations on post-label cut branches from the field (Fig. 1c).

### Tissue and sap water contents

In these potted saplings, the water remaining in the stem after centrifugation (denoted ‘tissue’ water from hereon) represented the majority of total stem water content (95% on average, or 0.37±0.01 g gDW^-1^ out of 0.39±0.01 g gDW^-1^). In the field, the tissue water of branches from adult trees also represented a large proportion of total stem water content (78 ± 5.6% on average, ranging from 70% to 92%, see also Fig. S1). Bulk stem water contents decreased mostly in spring and early summer (Fig. 2a). Tissue water content followed the same pattern and remained close to the range of values expected for the fiber saturation point of *F. sylvatica* wood (indicated by the hatched area in Fig. 2a) most of the summer, and recovered to spring levels only after leaf fall in November. Precipitation during spring and early summer 2018 was above average, and only a relatively mild drought was observed in late summer. This mild drought produced a progressive depletion of soil moisture in the upper soil (top 10 cm), while soil moisture content in deeper soil layers remained relatively stable, even at the peak of drought in September (Fig. 2b). Soil moisture at both depths started to increase to winter levels after the first autumn rains in October.

**Figure 2.**
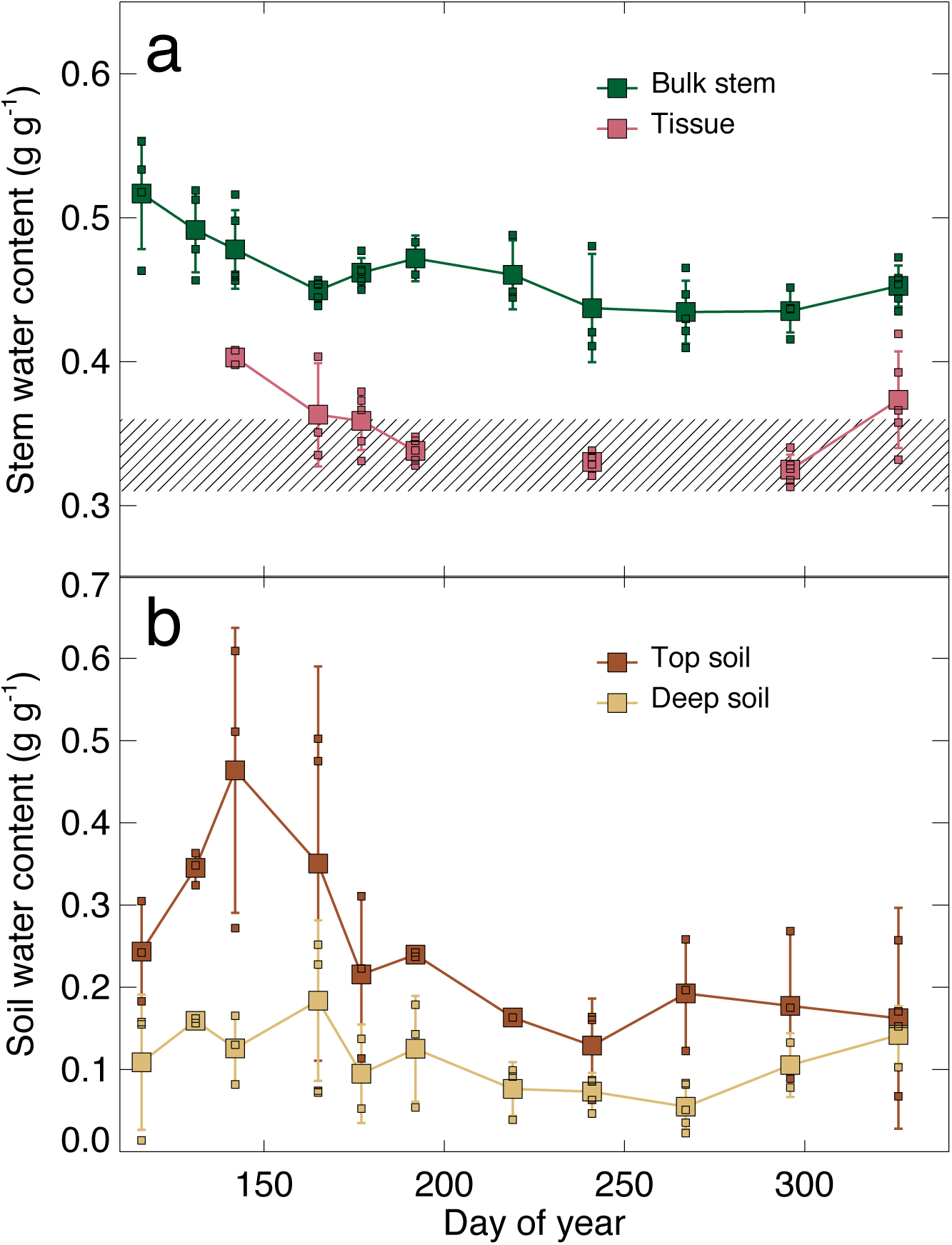
Temporal variations of gravimetric water contents during the 2018 growing season at the study site, a riparian forest in the SouthWest of France. (a) Bulk stem water and stem tissue water from *F. sylvatica* branches. (b) Deep and top soil water. In (a), the hatched area represents the expected range of wood fiber saturation point for *F. sylvatica* wood (Barkas, 1936; Berry & Roderick, 2005).

### Isotopic composition of sap and tissue water in the field

Over the growing season, there were notable differences in the isotopic composition of bulk stem water, sap water and tissue water of *F. sylvatica* branches. In the dual isotope space, sap water was always close to the LMWL, whereas the δ^2^H of tissue water was more negative than that of rain, stream water, ground water, or bulk soil water (Fig. 1e-f and Fig. 3). Bulk stem water exhibited intermediate δ^2^H values between sap and tissue water, suggesting that it represents a mixture of both pools. Sap water was always enriched in δ^2^H over bulk stem water (+16.6‰, *P* < 0.0001), whereas tissue water was always depleted in δ^2^H over bulk stem water, but to a lesser extent (−8.8‰, *P* < 0.0001). This latter result is expected by isotopic mass balance, because tissue water constitutes a larger proportion of total bulk water (Fig. 2a). Unlike δ^2^H, the δ^18^O of sap and tissue water was significantly more enriched than that of bulk stem water (+1.1‰, *P* < 0.0001 and +0.4‰, *P* < 0.001, respectively). However, when uncertainties were accounted for, the tissue-to-bulk water content ratio deduced from the combined δ^2^H and δ^18^O isotopic mass balance was always consistent with the tissue-to-bulk water content ratio measured gravimetrically (Fig. S1).

**Figure 3.**
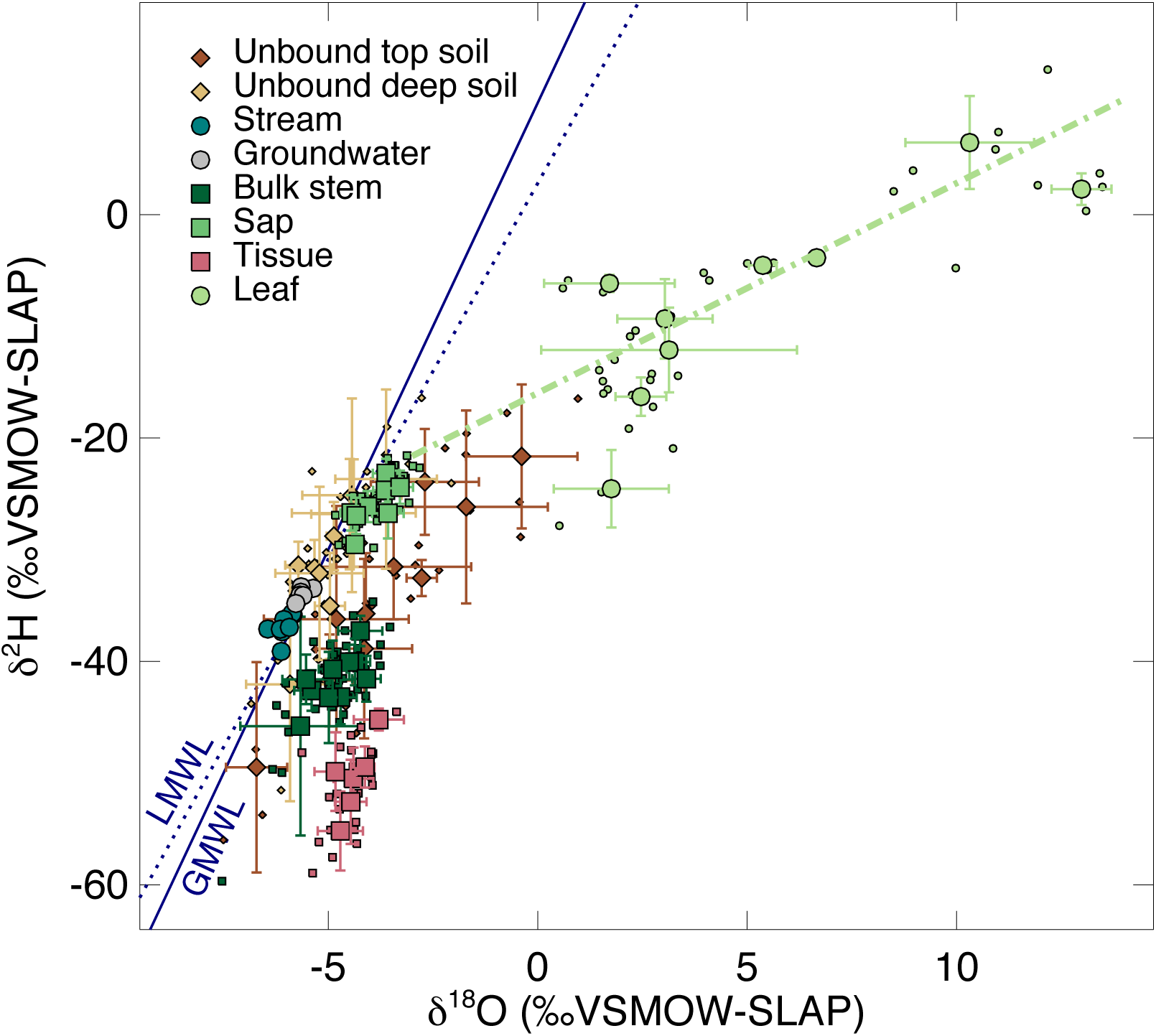
Dual isotope representation of the different water pools during the 2018 growing season at the study site, a riparian forest in the Southwest of France. Larger symbols represent daily means ± standard deviations for each sampling campaign, while smaller symbols are individual points (n = 3-5). LMWL: local meteoric water line. GMWL: global meteoric water line. The dot-dashed green line represents the ‘growing-season’ average leaf water evaporation line (see text).

### Plant water sources

The fact that tissue water content is always close to the fiber saturation point (Fig. 2a) is an indication that fiber-bound water must represent a large part of this stem water pool. It is therefore not surprising to observe strong hydrogen isotope effects on this water pool compared to other stem water pools, knowing that fiber-bound water is attached to cell walls through hydrogen bonds (Berry & Roderick, 2005). Similar surface effects have also been found for water films adsorbed onto mineral or organic surfaces (Chen *et al*., 2016; Lin & Horita, 2016; Lin *et al*., 2018). These studies imply that the isotopic composition of soil water accessible to plants should differ from that of bulk soil water. Chen and colleagues proposed empirical formulations to quantify the isotopic offset between bulk soil water and ‘unbound’ soil water, that is, soil water not adsorbed onto soil particles and more likely taken up by roots (see Methods). We applied this formulation to estimate the isotope composition of ‘unbound’ soil water from that of bulk soil water.

We found that unbound water in the deep soil layer had an isotopic composition that resembled that of rain, as it plotted on the LMWL (Fig. 1e-f and Fig. 3). A similar result had been observed in a soil pasture in Germany (Chen *et al*., 2016). Unbound topsoil water was more variable, following similar seasonal variations as bulk topsoil water. More interestingly, the soil evaporation line exhibited by bulk soil water (Fig. 1a-b) was still present for unbound soil water and now overlapped with sap water, while bulk stem water was taken even further away from it (Fig. 1e-f). Indeed, for each field campaign, bulk stem water δ^18^O and δ^2^H values rarely overlapped with those of ‘unbound’ soil water (Fig. 4). In contrast, for both δ^2^H and δ^18^O, sap water followed more closely variations in unbound water from deep soil layers. On a few rare occasions the isotopic composition of sap water became more enriched than top and deep soil water: at the beginning of July (day 192), after a few days of heavy rains (> 35 mm d^-1^) that depleted top soil water below that of deep soil water, and at the end of the growing season, just before and after leaf fall (days 296 and 326). In both situations, sap water δ^2^H and δ^18^O remained close to the values they had at the previous campaign (Fig. 4). In the former case this is consistent with the idea that the heavy rains in July displaced the relatively enriched unbound top soil water (Fig. 2b) to deeper intermediate horizons (located between the top and bottom sampling depths where it was still accessible to trees). This explanation is strongly supported by the enrichment of unbound water in the deepest soil layers in the subsequent sampling date. In contrast, the latter case is most likely caused by a reduction of transpiration accompanying leaf senescence that resulted in a slower isotopic turnover of sap water at the end of the season. At other dates over the growing season, our results indicate that trees mostly accessed water from deep soil layers (Fig. 4).

**Figure 4.**
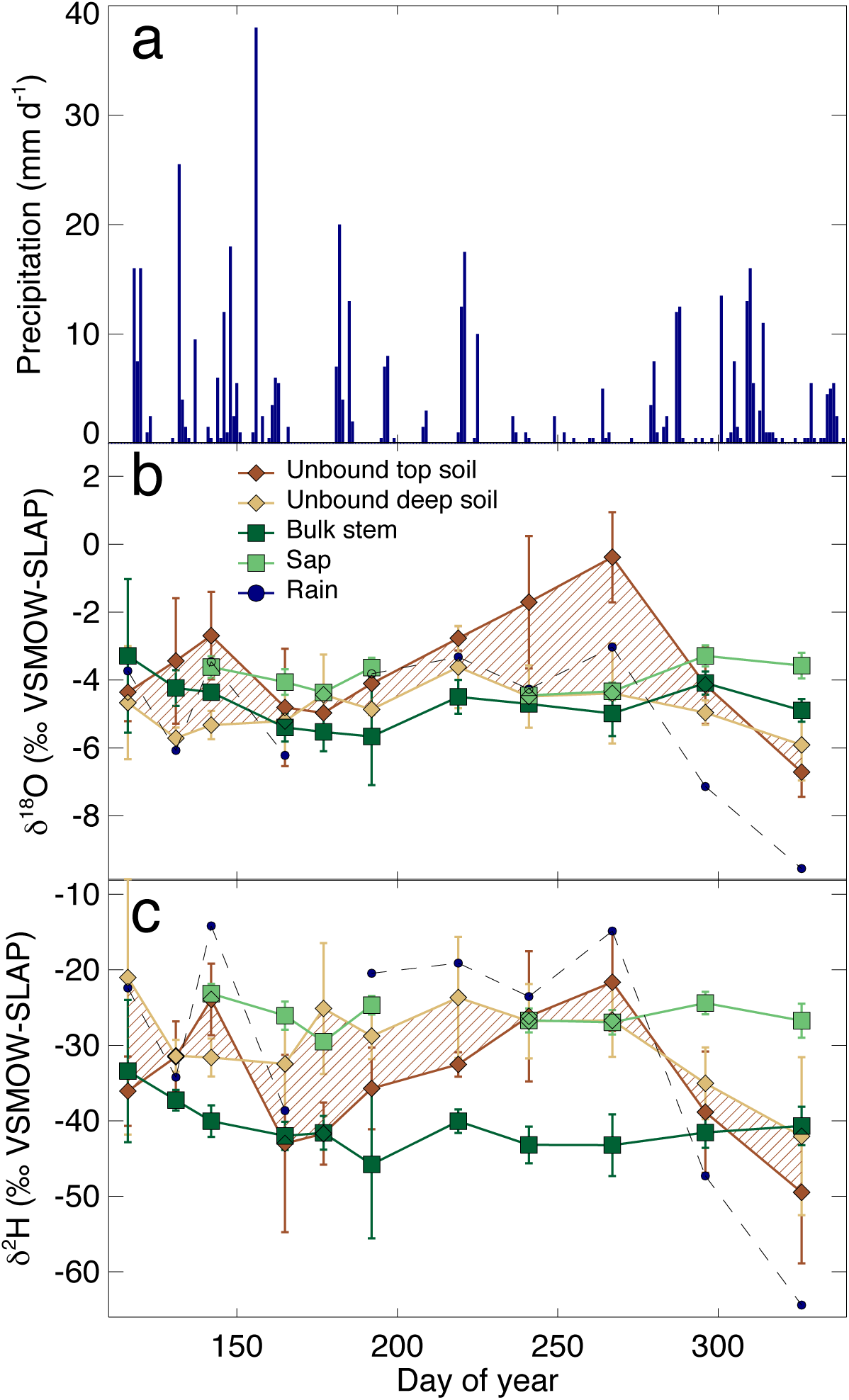
Seasonal variations of daily precipitation and ‘unbound’ top and deep soil water, bulk stem and ‘sap’ water, and rain water during the 2018 growing season. (a) Precipitation. (b) Oxygen isotope signals. (c) Hydrogen isotope signals. In (b-c), the hatched area is here to better visualize the range of values in the soil, taking top and deep soil as end members.

The isotopic stability of sap water and bulk stem water over the growing season is quite remarkable (Fig. 4) and is an indication that the depth of root water uptake did not suffer strong shifts over the growing season. Hence topsoil water, whose isotopic composition varies strongly over time and space, is unlikely a significant source of water for these riparian *F. sylvatica* trees. This isotopic stability of sap and bulk stem water pools also implies that the origin of leaf water should be temporally conservative as well. In these conditions, a linear regression in the dual isotope space of all the measured leaf water isotope data can be interpreted as a ‘growing season’ leaf evaporation line, whose intersection with the LMWL should correspond approximately to the source of leaf water. When performing such a regression, sap water is found as the most likely source of leaf water, in contrast to bulk stem water or tissue water that are too depleted in deuterium (Fig. 3).

## Discussion

### Which stem water pool is extracted with the cavitron technique?

The isotopic difference between sap water and bulk stem water (16.6‰ in δ^2^H) is of similar magnitude to the soil-stem isotopic offsets previously reported in glasshouse and field studies (Vargas *et al*., 2017; Barbeta *et al*., 2019, 2020; Poca *et al*., 2019). Thus our findings support the idea that isotopic compartmentalization within woody stems underpins these observed soil-stem isotopic offsets, complicating the identification of plant water sources. But can we be sure that water collected with the cavitron apparatus is more representative of the water being taken up by roots and transpired by leaves as opposed to bulk stem water obtained by cryogenic vacuum distillation? Three lines of evidence support this assertion.

Firstly, water extracted with the cavitron apparatus overlapped in the dual isotope space with unbound soil water from the deep horizons (Figs. 3-4), after accounting for isotopic fractionation processes occurring at the soil pore level. Unconfined soil water, also referred to as mobile soil water, is the soil water pool that is more likely to be accessed by roots (Bowling *et al*., 2017) and thus, its isotopic composition should be similar to that of sap water when it is available. Secondly, the growing-season average leaf water evaporative enrichment line had its origin in ‘cavitron-based’ sap water values, whereas both bulk stem water and tissue water were too depleted in ^2^H to be identified as the source of water for the leaves (Fig. 4). Thirdly, water extracted with the cavitron apparatus from potted saplings coincided well with irrigation water for both tap- and label-fed saplings (Fig. 1d). Small deviations between irrigation and ‘cavitron-based’ sap water could be explained by a slight evaporation enrichment of the cut stem after sampling (for tap-fed plants) and mixing of old and new sap water (for label-fed plants). In any case, the deviation from irrigation water was much smaller than the offset between sap and bulk stem water, demonstrating that the cavitron technique was suitable to characterize the isotopic composition of sap water separately from that of bulk stem water.

The fact that the water collected with the cavitron technique has an isotopic composition that matches that of sap water does not necessarily mean that the collected water is *only* sap water. It is generally thought that, during centrifugation, a cut branch would first release capillary water and water from open or large vessels, then from elastic living tissues and finally from smaller vessels and tracheids (Tyree & Yang, 1990). Following this principle, as most of the smaller conduits from *F. sylvatica* branches are embolized at -6 MPa (Stojnić et al., 2017), the water from living cells should have also been extracted alongside sap water from capillaries, vessels and tracheids. However if it were the case, we would expect to have co-extracted large amounts of organic compounds that would have affected the performance of the water isotope determination (Martín-Gómez *et al*., 2015), but this was not the case. In fact, recent studies using X-ray computed microtomography (microCT) imaging show that water held in fibers and living cells surrounding the conduits is released only after severe cavitation (e.g. Knipfer *et al*., 2019). This pattern has been observed in all microCT studies we are aware of, encompassing 13 woody species, on either intact plants (Cochard *et al*., 2015; Knipfer *et al*., 2015; Charrier *et al*., 2016; Choat *et al*., 2016; Li *et al*., 2020) or cut samples (Dalla-salda *et al*., 2014; Torres-Ruiz *et al*., 2015; Knipfer *et al*., 2019). This implies that the water extracted with the cavitron apparatus is mainly drawn from intercellular spaces and xylem conduits (our results on water extraction at two different rotation speeds indicating that they are hydraulically connected), leaving behind a large part of symplastic water from living cells (mainly parenchyma) and water within cell walls. The fact that the stem water content after centrifugation always remains close to the fiber saturation point (Fig. 2) is not in contradiction with the idea that symplastic water from living cells is still present. The fiber saturation point is usually measured on dead wood samples (i.e. without viable cells) but also under an atmosphere saturated in water vapor (usually >99% relative humidity, see Berry & Roderick, 2005). In our study, stem water contents were measured in room (and thus relatively dry) air. In this situation cell walls should not be saturated with water, while living cells should still contain some.

If a significant part of the ‘tissue’ water left in the sample after cavitation extraction is made up of water held in living parenchyma cells, surface isotopic effects during adsorption of water onto cellulosic fibers or other hydrophilic organic substances (Chen et al. 2016) may not be the only cause for the lower δ^2^H of ‘tissue’ water compared to sap water. In particular, the elastic fraction of symplastic water that continuously exchanges with xylem conduits may as well contribute to the overall isotopic depletion of tissue water compared to sap water, assuming that this water exchange through aquaporins (Pfautsch *et al*., 2015; Secchi *et al*., 2017) is a mass dependent fractionating process favoring the light isotope as previously postulated (Zhao *et al*., 2016; Poca *et al*., 2019). Additionally, symplastic water initially stored during cell formation, division and expansion could also be depleted in δ^2^H compared to sap water. In poplar, it has been observed that living fibers adjacent to the cambial region exhibit a high expression of aquaporins (Almeida-Rodriguez & Hacke, 2012). It is thus plausible that the origin of the lower δ^2^H in tissue water is caused during the channeling of water through aquaporins during cell formation. However, there is only indirect evidence that aquaporin-mediated transport may be a fractionating process (Mamonov *et al*., 2007). Further studies are required to test this hypothesis, which could be useful to understand and trace water movement within the xylem matrix. In contrast, surface isotope effects on water adsorbed to cellulosic fibers have already been demonstrated (Richard *et al*., 2007; Oerter *et al*., 2014; Chen *et al*., 2016) and are currently the most plausible explanation for water isotopic heterogeneity in woody stems.

Regardless of which process causes the isotopic depletion of tissue water compared to sap water, our labeling experiment demonstrated that it was possible to use the cavitron apparatus to extract sap water without isotopic fractionation (Fig. 1c-d), and without co-extracting large amounts of organic compounds. Therefore, the flow-rotor centrifuge-based extraction of sap water overcomes current methodological limitations in the application of stable isotopes to address several important questions in ecohydrology (Zhao *et al*., 2016; Barbeta *et al*., 2019, 2020; Oerter & Bowen, 2019).

### Implications of stem water isotopic heterogeneity for plant water source identification

The isotopic composition of bulk stem water has been extensively used to trace water fluxes in the soil-plant-atmosphere continuum and thus has been instrumental for elaborating hydrological and ecological theory of plant water use. Our results demonstrate that the δ^2^H of bulk stem water is different from the δ^2^H of water in the transpiration stream. In this respect, the analysis of the isotopic composition of sap may have led to significantly different conclusions in previous studies attributing plant water sources. For example, it is likely that isotope-based estimations of plant groundwater use (Barbeta & Peñuelas, 2017; Evaristo & McDonnell, 2017) would be slightly smaller if true sap water had been used instead of bulk stem water. Indeed, the soil-stem water offset may cause commonly applied isotope mixing models to underestimate the contribution of soil water to plant water use, whilst overestimating the contribution of isotopically-depleted ground water. Also, the isotopic differences between meteoric and runoff water and bulk stem water at the origin of the two water worlds hypothesis (Brooks *et al*., 2010; McDonnell, 2014) may have been much smaller if sap water had been used instead of bulk stem water, because its isotopic composition is much closer to that of the meteoric water line (Fig. 1 and 3) and thus to stream and ground water. The basis for the conceptual separation of belowground water pools into green water (accessed by plants) and blue water (contributing to groundwater and runoff) now requires revisiting. In particular, studies relying solely on δ^2^H to assess the origin of plant water (e.g. Allen *et al*., 2019) may be especially sensitive to the effects of within-plant isotopic heterogeneities.

It is only recently that soil-stem water isotopic offsets have been reported in plants that are not halophytes or xerophytes (Vargas *et al*., 2017; Barbeta *et al*., 2019; Oerter & Bowen, 2019; Barbeta *et al*., 2020). In a previous study on *F. sylvatica* saplings, it was shown that such isotopic offsets could not be attributed to isotopic fractionation during root water uptake (Barbeta *et al*., 2020). Instead, and consistent with previous studies (White *et al*., 1985; Yakir *et al*., 1994; Zhao *et al*., 2016), it was suspected that within-stem isotopic heterogeneities were responsible for the isotopic differences between soil water and bulk stem water. Here, by applying for the first time ‘cavitron-based’ sap water extractions, isotopic differences between sap water and other xylem tissues is confirmed in both controlled and field settings. This key result should lay the foundations for the development of new paradigms in isotope ecohydrology.

## Materials and Methods

### Field site

The study site was a mixed riparian forest on the karstic canyon formed by the Ciron, a tributary of the Garonne river in SW France (44°23 N, 0°18 W, 60 m a.s.l.). Soil texture ranges from coarse sand at the surface to loamy coarse sand in the deeper horizons, where the presence of limestone rocks weathered to various degrees creates a distinguishable carbonate-rich C horizon (Table S1). This riparian forest is dominated by deciduous species including *F. sylvatica* and *Q. robur*. The site has a temperate oceanic climate (Cfb in the Köppen-Geiger classification). Daily meteorological data was available from a weather station located at about 20 km from the studied site, and long-term (1897-present) monthly temperature and precipitation data was also available from another weather station located 16 km away from the studied area. Over the period 1897-2015, the mean annual temperature was 12.9°C and the mean annual precipitation was 813 mm y^-1^, distributed rather evenly over the seasons.

### Stem, soil and water sampling

For the present study, we selected one of the plots sampled in a previous study (Barbeta *et al*., 2019), in which a more detailed description of the plots is available. In 2018, we conducted sampling campaigns each month over the entire growing season. Five dominant *F. sylvatica* trees were chosen for collecting different stem water pools. From long (>1m length), relatively straight branches, a sub-section, a few cm in length, was cut and used for bulk stem water. After removing the bark and phloem in the field, the sub-section was immediately placed in an air-tight Exetainer® sealed with Parafilm® and kept in a cool box until storage in the lab at 4°C. Leaves belonging to that sub-branch were also collected. The central stem of each branch was re-cut to a segment of *ca*. 50 cm in length and, without peeling off the bark, the open cuts were covered with Parafilm®. These longer stem segments were sealed in plastic bags and also stored at 4°C once in the lab to minimize stem evaporation. Sap water was always extracted from those longer stems within 24 hours from their collection in the field. On fewer occasions, the same sampling procedure was used on *Q. robur* and *P. pinaster* branches, to test the novel extraction method (see below). Additionally, for each species, a subset of five of these stem samples have been flushed with isotopically labelled water (δ^18^O = 2.9± 0.2 ‰; δ^2^H = 8.5 ± 0.6‰) during at least 2h at a pressure of 1.8bar for vessel-bearing species (*F. sylvatica* and *Q. robur*) and during 18h at 0.1 bar for tracheid-bearing species (*P. pinaster*).

In addition to plant material, three soil cores amongst the sampled trees were extracted with a soil auger. Each soil core was split into topsoil (0-10 cm) and deep soil (from 70-80 to 110-120 cm depending on the depth of the bedrock). Soil samples were placed in 20 mL vials with positive insert screw-top caps, sealed with Parafilm® and kept in a cool box until they were stored in the lab at 4°C.

Samples of stream water, groundwater (from a well located ca. 50 m from the river) and fog and rain water (from collectors installed in a small open area about 100 m away from the sampling plot) were also collected in each campaign. Details on the rain collector can be found in a previous paper (Barbeta *et al*., 2019).

### Experiment on potted plants

In order to further test the proposed methodology to extract sap water from woody stems, we also grew potted saplings of *Fagus sylvatica* and *Quercus robur*. During 18 days, 7-10L pots with 1-1.5 m height saplings were placed into a climate-controlled growth chamber with 16-8 h day-night cycle (photosynthetically active radiation: 700 µmol m^-2^ s^-1^; day temperature; 24°C; night temperature; 22°C; relative humidity: 40-50%). Each pot was placed on automated balances to monitor water losses and irrigated daily with tap water (δ^18^O = - 38.19±0.32‰; δ^2^H = -6.43±0.10‰) until field capacity. We covered the soil with aluminum foil to prevent soil evaporation and isotopic heterogeneity with depth. During the last three days of the experiment, we switched irrigation water from tap to label water (δ^18^O = 2.91±0.01‰; δ^2^H = 5.72±0.08‰) for 3 of the 7 saplings (label-fed plants), and increased the irrigation rate (3L d^-1^) to ensure near-complete replacement of soil water with label water. The other 4 pots were irrigated at the same rate but with tap water (tap-fed plants). Transpiration rate ranged from 0.18 to 0.32 L d^-1^ and was similar in both tap-fed and label-fed plants. On the last day of the irrigation period, at midday, trees were harvested and the main stem was cut and prepared for immediate centrifugation and cryogenic extraction, following the same procedure as that used for branches from the field (see above). Soil gravimetric water content was also measured and averaged around 0.7 g g^-1^ with no differences between treatments or species.

### Water extraction methods

Bulk stem and soil water was extracted from fresh samples using a vacuum line (<10 Pa). Once in the vacuum line, samples were heated to 80°C for 2-3h and the evaporated vapors were collected in a cold trap. A detailed description of the design and methodology can be found in our previous studies (Barbeta *et al*., 2019, 2020). Stem ‘tissue’ water was extracted using the same methodology from sub-sections of the longer branches used to extract sap water (see below). Gravimetric water content and extraction yields were assessed for each soil and plant sample by weighing the sample before and after water extraction. We also checked that the water extraction had been completed by oven drying all samples at 105°C for 24h and re-weighing them.

Sap water was extracted using a flow-rotor centrifuge originally designed to study the hydraulic embolism resistance of woody stems (Cochard, 2002) and equipped with custom-made water collectors. These collectors were made of a plastic reservoir fitted with a watertight resin lid, enabling the insertion of the woody stem in the reservoir whilst preventing the extracted water from escaping the bottom of the reservoir once spinning stopped. Branches where spun first at 3130 rpm for 120 s, which corresponds to a minimum negative pressure in the middle of the branch of about -2 MPa. Rotation was then stopped, and the liquid extracted from the branch was collected from both upstream and downstream reservoirs. The reservoirs and the sample were then placed back in the rotor, and spun at 5280 rpm for another 120 s, corresponding to a minimum pressure of -6 MPa. Rotation was stopped again and the liquid extracted from the branch was collected similarly. A section about 10 cm long was then cut from the centre of the branch to extract the ‘tissue’ water remaining in the sample using the cryogenic vacuum extraction line described above.

As with any other sampling technique to measure the isotopic composition of water, the evaporation of water in the samples must be avoided, as it strongly modifies the isotopic composition of water in the samples. Based on our data from the labelling experiments (Fig. 1 c-d), it seems that evaporation during centrifugation is negligible. Notably, in an experimental setup similar to the one used here to extract sap water, it has been shown that evaporation of the collected water operates at a rate of around 0.5 mg min^-1^ when spinning at 6000 rpm (Peng *et al*., 2019). In our case, this corresponds to about 0.1% of the water volume collected after 2 minutes of spinning (typically 1 mL). Therefore, evaporation during spinning should affect only marginally the isotopic composition of the cavitron-extracted water. On the other hand, we did not quantify potential evaporation from samples collected in the field, where the isotopic composition of source water was uncertain. Although care was taken to prevent evaporation during field sample collection and transportation (see Methods), this step is probably the most susceptible to induce evaporative enrichment of the sampled water. Nevertheless, our results from both field and glasshouse experiments demonstrate that the cavitron technique provides a much closer estimate of the isotopic composition of sap (and thus of the plant water), compared to cryogenic extraction.

### Water isotope analyses

The isotopic composition (δ^2^H and δ^18^O) of the different waters were measured with an off-axis integrated cavity optical spectrometer (TIWA-45EP, Los Gatos Research, USA) coupled to an auto-sampler. Details on the precision of the instrument, calibration and post-correction procedures, notably to account for the presence of organic compounds, can be found in previous studies (Barbeta *et al*., 2019, 2020). All isotopic data reported here are expressed on the VSMOW-SLAP scale.

### Estimating the isotope composition of unbound soil water

Isotopic offsets between bulk and unbound soil water estimated using the empirical formulations proposed by Chen and colleagues for organic and mineral surfaces (Chen *et al*., 2016). Because the organic fraction of our soil samples was negligible, we simplified these formulations to:

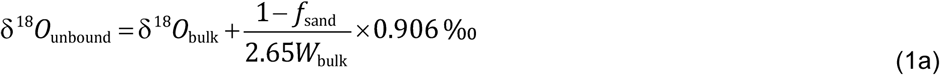

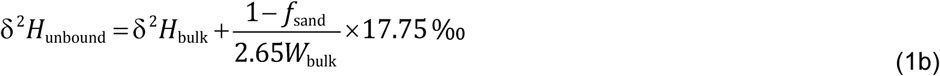

where *f*_sand_ and *W*_bulk_ represent the sand fraction and gravimetric water content of the soil sample, respectively. The sand fraction was subtracted because only finer minerals are considered to create the isotopic offset due to their greater ability to attract large amounts of adsorbed water (Chen *et al*., 2016). An average sand fraction of 0.92 was used based on texture analysis of the different soil horizons (Table S1).

### Data analyses

Statistical differences between the isotopic composition of bulk stem water, sap water and tissue water from field samples, as well as between those of labelled tissue and vessel water were assessed with Generalized Linear Models from the package *lmer* (Bates *et al*., 2015) in *R* (R Core Team, 2019). The same models were also used to assess statistical differences between bulk and tissue water gravimetric water contents and the isotopic composition of water extracted at -2 and -6 MPa. These models allow us to set random factors such as date of sampling and/or tree individual, when necessary.

## Acknowledgments

Many thanks to Anne-Isabelle Gravel, Gaëlle Capdeville and Sylvain Delzon for assistance at the Cavitron facility and to Laura Clavé and Kenza Bakouri for assistance in the field. This study received funding from the EC2CO/BIOHEFECT program (CNRS, France), the French national research agency (projects Leafshed and Hydrobeech within the Cluster of Excellence COTE with grant agreement ANR-10-LABX-45; project ORCA with grant agreement ANR-13-BS06-0005-01), the European Research Council (ERC) under the EU Seventh Framework Program (FP7/2007-2013, with grant agreement no. 338264, awarded to L.W.) and the Aquitaine Region (project Athene with grant agreement 2016-1R20301-00007218). A.B. and P.M. also acknowledge funding from IdEx Bordeaux postdoctoral fellowships from the Université de Bordeaux (ANR-10-IDEX-03-02).

## Figures and Tables

**Figure S1.**
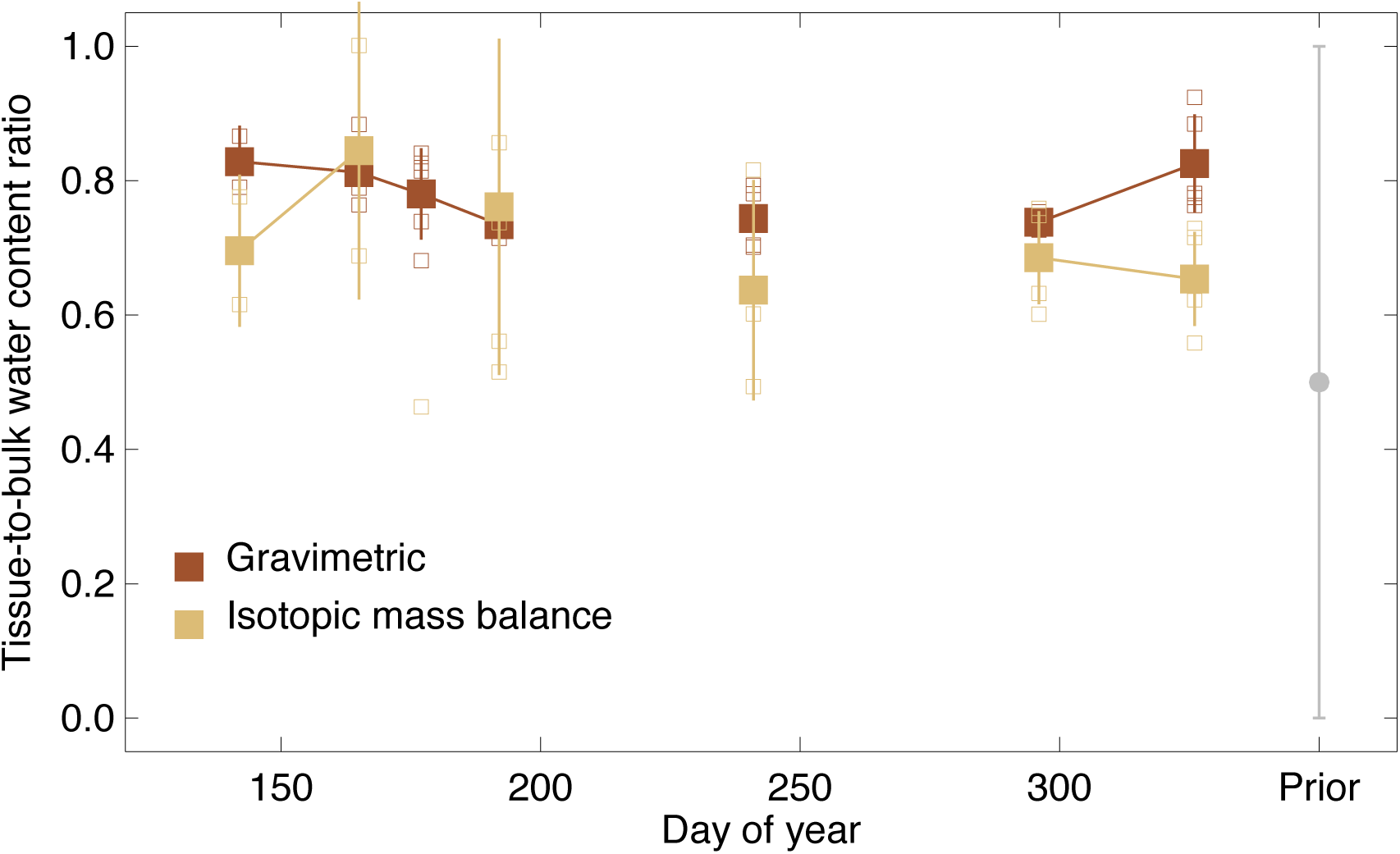
Temporal variations of tissue-to-bulk water content ratios during the 2018 growing season, estimated either by gravimetric measurements or by isotopic data and isotopic mass balance. The latter was estimated for each individual branch using a Bayesian optimization approach that takes into account the isotopic mass balance for the two water isotopes, the uncertainty on the isotopic data and a prior value of the ratio of 0.5 ± 0.5. Larger symbols represent the mean ± standard deviation (n =3-5).

**Table S1.**
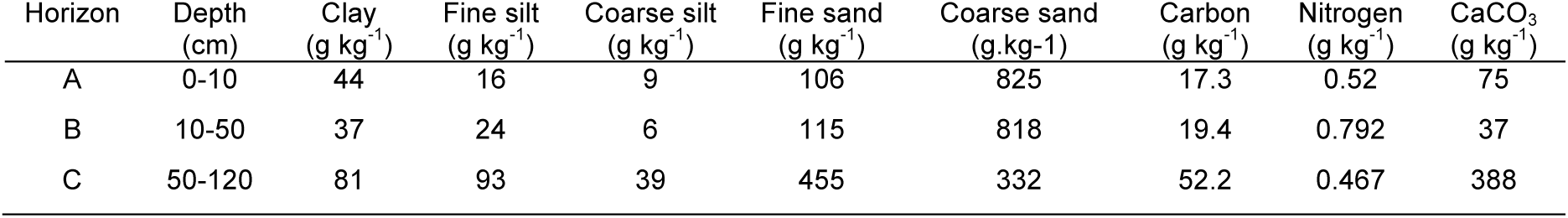
Soil properties at the study site, the Ciron river gorges in SW France.

| Horizon | Depth<br>(cm) | Clay<br>(g kg<sup>-1</sup>) | Fine silt<br>(g kg<sup>-1</sup>) | Coarse silt<br>(g kg<sup>-1</sup>) | Fine sand<br>(g kg<sup>-1</sup>) | Coarse sand<br>(g.kg<sup>-1</sup>) | Carbon<br>(g kg<sup>-1</sup>) | Nitrogen<br>(g kg<sup>-1</sup>) | CaCO<sub>3</sub><br>(g kg<sup>-1</sup>) |
| --- | --- | --- | --- | --- | --- | --- | --- | --- | --- |
| A | 0-10 | 44 | 16 | 9 | 106 | 825 | 17.3 | 0.52 | 75 |
| B | 10-50 | 37 | 24 | 6 | 115 | 818 | 19.4 | 0.792 | 37 |
| C | 50-120 | 81 | 93 | 39 | 455 | 332 | 52.2 | 0.467 | 388 |

